# Linking White Matter Integrity to Recognition Memory Speed: Fixel-Based and Fornix Analyses in Young to Middle Adulthood

**DOI:** 10.1101/2025.07.17.665387

**Authors:** Melissa Elder, Timothy M. Ellmore

## Abstract

Age-related differences in white matter structure are increasingly understood as nonlinear, regionally specific, and behaviorally relevant. Using whole-brain fixel-based analysis (FBA), we examined how recognition memory speed relates to micro- and macrostructural white matter properties across a sample spanning young to middle adulthood. Slower response times were associated with higher fiber density (FD) in left frontoparietal tracts, nonlinear increases in fiber cross-section (log(FC)) in the anterior corpus callosum, and elevated combined fiber density and cross-section (FDC) in posterior callosal and cingulum pathways. These associations were most pronounced among individuals in the later decades of this age range, suggesting that white matter morphology reflects both extended maturation and emerging age-related decline. In a separate, hypothesis-driven analysis, we applied deterministic tractography to reconstruct the fornix and extracted mean fractional anisotropy (FA) along its length. Greater fornix FA and younger age together explained 34% of the variance in retrieval speed. These findings highlight regionally distinct structural contributions to memory performance and support lifespan models emphasizing individual variability and neuroplasticity in white matter development. This integrative approach underscores the value of combining whole-brain and tract-specific analyses to advance our understanding of white matter contributions to cognitive aging.

**Highlights:**

- Slower RT linked to higher fiber density in left frontoparietal white matter
- Slower RT associated with nonlinear increases in anterior corpus callosum log(FC)
- FDC increased in posterior callosal and cingulum tracts in slower older adults
- Findings support prolonged maturation and regionally specific age-related decline
- Fornix FA and age explained 34% of variance in memory retrieval speed

## Graphical Abstract

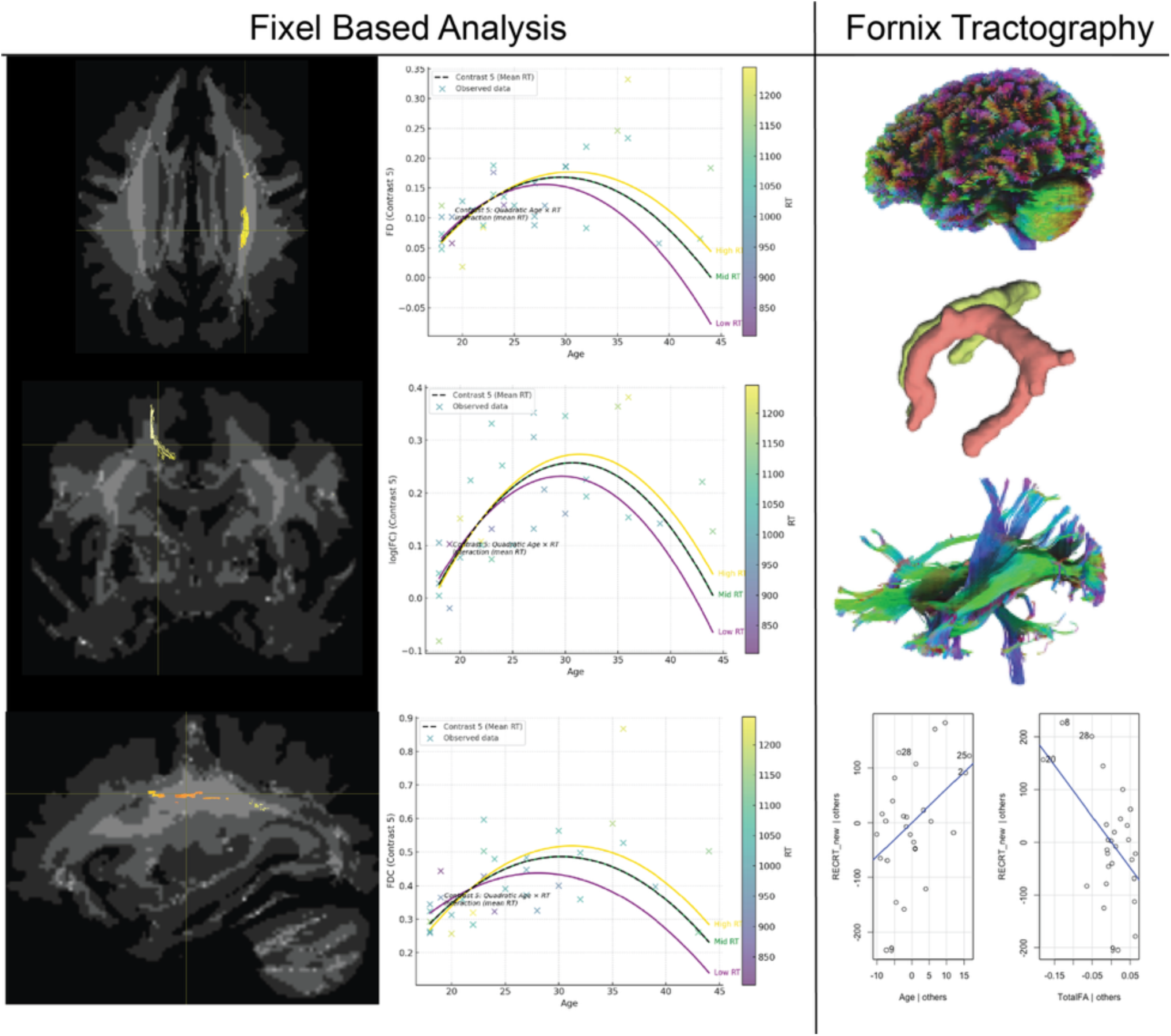

## Introduction

The fornix is a major white matter tract that plays a crucial role in memory processing by connecting the hippocampus to the mammillary bodies, anterior thalamic nuclei, and other subcortical structures (Aggleton et al., 2000). As a key component of the extended hippocampal-diencephalic circuit, the fornix is critically involved in episodic memory and long-term recollection (Tsivilis et al., 2008). Lesion studies have demonstrated that damage to the fornix produces profound anterograde amnesia, with disproportionate deficits in recall compared to recognition memory (Aggleton et al., 2000; Park et al., 2000). Advances in neuroimaging, particularly diffusion tensor imaging (DTI), have provided new insights into how fornix microstructure relates to various types of memory function, including episodic memory, visual recognition memory, and working memory (Mielke et al., 2012; Rudebeck et al., 2009).

Episodic memory is one of the primary cognitive functions affected by fornix disruption. Studies of patients with bilateral fornix lesions, such as those resulting from colloid cyst surgery, reveal profound impairments in free recall tasks but relatively preserved recognition memory (Aggleton et al., 2000; Tsivilis et al., 2008). These findings are supported by DTI research demonstrating that fornix integrity, as measured by fractional anisotropy (FA), is a strong predictor of episodic recall performance in both healthy individuals and those with memory disorders (Mielke et al., 2012). Furthermore, fornix atrophy and reduced FA have been linked to accelerated memory decline in individuals with mild cognitive impairment (MCI), highlighting the fornix as an early biomarker of Alzheimer’s disease (Mielke et al., 2012).

While the fornix is essential for episodic memory, its role in recognition memory is more nuanced. Recognition memory is often conceptualized as involving two components: recollection and familiarity (Yonelinas, 2002) Recollection involves retrieving detailed contextual information, whereas familiarity is a more automatic sense of knowing. Studies suggest that the fornix is selectively involved in recollection-based recognition, as individuals with fornix lesions exhibit deficits in recollection but retain relatively intact familiarity-based recognition (Rudebeck et al., 2009). DTI studies confirm that fornix FA is significantly correlated with recollective recognition but not with simple familiarity judgments, reinforcing the idea that the fornix primarily supports context-rich memory retrieval (Rudebeck et al., 2009).

In contrast to its clear role in episodic and recollective recognition memory, the fornix appears to play a minimal role in working memory. Patients with isolated fornix damage typically exhibit intact working memory, as assessed by tasks such as digit span or short-delay recall (Gaffan et al., 1991; Zahr et al., 2009). However, some DTI studies have reported weak correlations between fornix integrity and working memory performance in aging populations, suggesting that fornix- related changes may influence general cognitive function rather than serving as a direct neural substrate for working memory (Zahr et al., 2009). This distinction underscores the specificity of the fornix’s contribution to long-term memory processes rather than transient information maintenance.

Reaction time studies suggest that recognition memory is neither purely recall-like nor purely familiarity-based, but rather involves two separable processes. Fast reaction times (RTs) are typically associated with familiarity-based recognition, which is thought to be an automatic and heuristic-driven process, whereas slower RTs suggest a more effortful, recollection-based retrieval process (Mickes et al., 2010). Studies using the Remember/Know paradigm have demonstrated that "know" responses (familiarity) are associated with faster RTs than "remember" responses (recollection), further supporting a dual-process model of recognition memory (Yonelinas & Jacoby, 1995). Additionally, neuroimaging research has linked fast recognition decisions to perirhinal cortex activity, a region implicated in familiarity processing, while slower, more deliberate recollection judgments recruit hippocampal and fornix pathways (Diana et al., 2006; Rugg & Curran, 2007). These findings underscore the need to examine how fornix integrity influences processing speed in different types of memory retrieval.

Given the complexity of memory retrieval and its reliance on distributed white matter networks, we first employed a whole-brain fixel-based analysis (FBA) to identify age- and performance-related variations in fiber density (FD), fiber cross-section (FC), and their combined metric (FDC). Unlike conventional diffusion tensor imaging (DTI), FBA allows for improved anatomical specificity in regions with crossing fibers and enables distinction between microstructural and macrostructural white matter properties. This approach provided a data-driven means of detecting tract-specific morphological changes related to recognition memory performance and aging, particularly in large association pathways involved in attentional and integrative processes.

To complement the exploratory whole-brain analysis, we conducted a hypothesis-driven tractography analysis of the fornix, a small, anatomically specialized white matter pathway known to support episodic and recollective memory processes. Prior work has linked fornix microstructure, particularly fractional anisotropy (FA), to episodic recall and context-rich recognition (Aggleton et al., 2000; Mielke et al., 2012; Rudebeck et al., 2009), while largely sparing familiarity-based and working memory functions (Zahr et al., 2009). Because recognition memory can be decomposed into fast, familiarity-based responses and slower, recollection-based judgments (Mickes et al., 2010; Yonelinas & Jacoby, 1995), we examined reaction time as a continuous behavioral index of retrieval efficiency. Our primary goal was to determine whether individual differences in fornix integrity predict recognition memory speed across adulthood and whether these effects are distinct from broader, white matter maturation detectable via whole-brain FBA. By integrating both exploratory whole-brain and region-specific hypothesis-driven methods, this study aims to clarify how distinct structural mechanisms contribute to age-related differences in memory retrieval speed and to identify whether microstructural preservation of the fornix plays a unique role in supporting efficient, recollection-based recognition.

## Methods

### Participants

Thirty-two healthy adults (ages 18-44 years) were recruited from City College, the CUNY Graduate Center, and the broader New York City community. Recruitment was conducted through flyers posted on campus and web postings via the City College of New York SONA online experimental scheduling system. The study was conducted under a protocol approved by the Institutional Review Board of The City University of New York Human Research Protection Program (CUNY HRPP IRB). All methods adhered to the guidelines and regulations of the CUNY HRPP IRB committee. Each participant provided written informed consent before participation.

All participants had normal or corrected-to-normal vision and reported no history of neurological or psychiatric disorders. Due to a technical failure that prevented response recording, data from five participants (two from Group A and three from Group B) were excluded, resulting in a final sample of 27 participants (14 female, 12 male, 1 non-binary). The participant identifying as non-binary was excluded from sex-specific analyses. The mean age of the final sample was 26 years (SD = 8). Exclusion criteria included claustrophobia, the presence of non-MRI-compatible metals (including some tattoos), and severe visual impairment requiring corrective lenses beyond ±7 diopters. Participants were compensated at a rate of $20 per hour for up to two hours of participation.

### Experimental Design

The experiment was conducted in a single 2-hour session inside a 3 Tesla MRI scanner. The design included alternating rest and memory tasks. The session began with the acquisition of field maps, followed by a 5-minute eyes-open resting-state fMRI scan, during which participants fixated on a central crosshair. After this initial rest scan, participants completed three consecutive 12- minute working memory task runs, with 2-minute breaks between each run for additional field map acquisition.

After the final working memory run, participants rested with their eyes closed while a high-resolution T1-weighted anatomical scan was acquired. This was followed by a second 5-minute resting-state fMRI scan, a high-resolution T2-weighted anatomical scan, and another set of field maps. The collection of these anatomical and resting-state fMRI data took approximately 20 minutes.

Next, participants completed a long-term recognition task during a single 10-minute fMRI run. Finally, the session concluded with two diffusion MRI acquisitions.

### Working Memory (WM) Task

Each WM run consisted of 52 trials of a Sternberg working memory task, each following a structured sequence designed to assess short-term memory encoding, maintenance, and retrieval processes. Each trial began with an encoding phase, during which three stimuli were presented sequentially, each for 1400 milliseconds (ms). This was followed by a 5000 ms delay period, during which no stimuli were shown, allowing for short-term memory retention. After the delay, a probe stimulus appeared for 1400 ms, requiring the participant to determine whether it was part of the previously encoded set. Immediately following the probe, a scrambled stimulus was presented for 2000 ms, serving as a perceptual mask and signaling the end of the trial. To temporally decorrelate the onsets of successive trials, a variable inter-trial interval (jitter) was introduced between the end of the scrambled stimulus and the beginning of the next encoding phase. The average jitter duration was 2298 ms. Each run contained 28 positive probes (where the probe matched an item from the encoding set) and 24 negative probes (where the probe did not match), resulting in a slightly unbalanced probe distribution (∼54% positive). A strictly equal split was not used in order to maintain flexibility in trial randomization and avoid introducing predictable patterns, while still preserving a near-equal ratio to support unbiased comparisons between trial types. This full sequence engaged encoding, maintenance, and retrieval processes central to working memory function.

### Recognition Memory Task

The long-term recognition task was administered as a single fMRI run approximately 20 minutes after the final working memory (WM) run. Participants were presented with a continuous stream of 340 stimuli, consisting of 180 positive probes (previously seen during the WM runs), 60 negative probes (novel, unseen stimuli), and 100 scrambled stimuli (perceptual controls). Each stimulus was displayed for 1000 milliseconds (ms) and was followed by a variable inter-stimulus interval (ISI), with an average ISI of approximately 1045 ms. The stimuli were randomly intermixed, and participants were instructed to indicate whether each item had been presented earlier in the experiment (positive), was new (negative), or was a scrambled non-item. This design allowed for the assessment of long-term memory recognition and discrimination between previously encountered and novel stimuli, while controlling for low-level visual processing using scrambled images.

### Stimuli

Validated pictures of natural scenes and objects were used for both the working memory and recognition memory tasks. One set of participants (Group A) were presented with indoor and outdoor scene images (Xiao et al., 2010), while another set (Group B) viewed both scene images and images of common objects (Jiang et al., 2022). Behavioral and diffusion data from both groups were combined for statistical analysis.

### Behavioral Analysis

Performance on the working memory (WM) and recognition memory tasks was quantified as the percentage of correctly identified images, defined as the proportion of old images correctly recognized and new images correctly rejected. This combined measure was reported as percentage correct.

For the WM task, mean reaction time (RT) and the average RT difference between the first and third runs were calculated to track improvement in RT from across the entire set of WM trials. For the recognition task, mean RTs were calculated separately for old and new images.

### Magnetic Resonance Imaging

MRI data were acquired using a 3T Siemens Prisma scanner. Whole-brain T1-weighted structural images were collected using an MPRAGE sequence with the following parameters: 1.0 mm isotropic resolution, repetition time (TR) = 2400 ms, echo time (TE) = 2.15 ms, inversion time (TI) = 1000 ms, flip angle = 8°, bandwidth = 220 Hz per pixel, and a 256 × 240 sagittal acquisition matrix.

Diffusion-weighted imaging (DWI) data were acquired using two high-angular-resolution spin-echo EPI scans, varying in the number of diffusion directions and phase encoding direction. Acquisition parameters included: 1.5 mm isotropic resolution, TR = 3230 ms, TE = 89.20 ms, flip angle = 78°, bandwidth = 1700 Hz per pixel, echo spacing = 0.69 ms, partial Fourier factor = 6/8, no iPAT, and a multi-band slice acceleration factor of 4. The scans included 92 directions with anterior-to-posterior (AP) phase encoding and 92 directions with posterior-to-anterior (PA) phase encoding. Diffusion volumes were acquired at b-values of 0, 1495, and 2995 s/mm². Preprocessing was performed using QSIPrep 0.15.4 (Cieslak et al., 2021).

### Anatomical Data Preprocessing

The T1-weighted (T1w) image was corrected for intensity non-uniformity using N4BiasFieldCorrection (ANTs 2.3.1; (Tustison et al., 2010)) and designated as the T1w reference for subsequent processing. Skull stripping was performed using antsBrainExtraction.sh (ANTs 2.3.1) with the OASIS template as the target.

Spatial normalization to the ICBM 152 Nonlinear Asymmetrical template (version 2009c) (Fonov et al., 2009) was conducted via nonlinear registration using antsRegistration (ANTs 2.3.1; (Avants et al., 2008)), applying brain-extracted versions of both the T1w image and the template.

Brain tissue segmentation of cerebrospinal fluid (CSF), white matter (WM), and gray matter (GM) was performed using FAST (FSL 6.0.5.1; (Zhang et al., 2001)) on the skull-stripped T1w image.

### Diffusion Data Preprocessing

Diffusion-weighted images (DWIs) were grouped into two phase encoding polarity groups. Images with b-values below 100 s/mm² were treated as b = 0 images. MP-PCA denoising was applied using MRtrix3’s dwidenoise (Veraart et al., 2016) with a 5-voxel window. B1 field inhomogeneity was then corrected using dwibiascorrect (MRtrix3) with the N4 algorithm (Tustison et al., 2010). Following bias correction, the mean intensity of each DWI series was adjusted to match across scanning sequences. Both phase encoding groups were then merged into a single file for processing with FSL workflows.

Head motion and eddy current distortions were corrected using FSL eddy (version 6.0.5.1:57b01774; (Andersson & Sotiropoulos, 2016)) with a q-space smoothing factor of 10, five iterations, and 1000 voxels used for hyperparameter estimation. A linear first- and second-level model was applied to characterize eddy current-related spatial distortions, and q-space coordinates were forcefully assigned to shells. Field offset and subject motion were estimated separately, and shells were aligned post-eddy. Eddy’s outlier replacement (Andersson et al., 2016) was enabled, grouping data by slice and replacing values from slices containing at least 250 intracerebral voxels when deviations exceeded four standard deviations.

Reversed phase-encode blips were used to acquire multiple DWI series with opposite distortion patterns. b = 0 images were extracted from each series to estimate the susceptibility-induced off-resonance field using a method similar to (Andersson et al., 2003). The estimated fieldmaps were incorporated into the eddy current and motion correction step, with final interpolation performed using the jac method.

Several confounding time-series were calculated from the preprocessed DWI data, including framewise displacement (FD) (as implemented in Nipype following (Power et al., 2014)). Head motion estimates from the correction step were added to the confounds file, along with slicewise cross-correlation measures. DWIs were then resampled to ACPC space with 1.5 mm isotropic voxels.

Internal operations of QSIPrep utilized Nilearn 0.9.1 (Abraham et al., 2014) and Dipy (Garyfallidis et al., 2014).

### Fixel-Bases Analyses

We conducted a whole-brain fixel-based analysis (FBA) (Raffelt et al., 2017) to examine how recognition memory performance and age relate to white matter structure. A general linear model was constructed with five regressors: an intercept, z-scored age, z-scored recognition reaction time (RT), a quadratic term for z-scored age (zAge²), and an interaction term between zAge and zRT. Twelve planned contrasts were tested to capture linear, interaction, and nonlinear effects. Linear effects were modeled by the main effects of age [0 1 0 0 0] and RT [0 0 1 0 0], as well as their negative counterparts. Joint additive effects were assessed using contrasts for age + RT [0 1 1 0 0], and age – RT [0 1 –1 0 0]. The interaction between age and RT was explicitly modeled using the zAge × zRT term [0 0 0 0 ±1]. Quadratic (nonlinear) age effects were captured by the zAge² term [0 0 0 ±1 0]. All continuous predictors were z-scored to reduce collinearity. Statistical inference was performed using nonparametric permutation testing (5,000 permutations) with family-wise error (FWE) correction. Each contrast was evaluated separately for three fixel-wise metrics: fiber density (FD), log-transformed fiber cross-section (log(FC)), and their combined product, fiber density and cross-section (FDC).

### Tractography

Tractography was performed using DSI Studio (Yeh et al., 2021) on preprocessed diffusion data. A multishell diffusion scheme was used with b-values of 1495 and 2995 s/mm², 92 diffusion sampling directions per shell, an in-plane resolution of 1.5 mm, and a slice thickness of 1.5 mm. The accuracy of the b-table orientation was verified by comparing fiber orientations with a population-averaged template (Yeh et al., 2018), and the b-table was flipped by 0.012fy. Restricted diffusion imaging quantified restricted diffusion (Yeh et al., 2017), and diffusion data were reconstructed using generalized q-sampling imaging (GQI) (Yeh et al., 2010)with a diffusion sampling length ratio of 1.25. Tensor metrics were calculated using only DWIs with b-values below 1750 s/mm².

A deterministic fiber tracking algorithm (Yeh et al., 2013) with augmented tracking strategies (Yeh, 2020)was applied to each region of interest (ROI). The HCP842 Tractography Atlas was used to define ROIs in the left fornix (70,81,57; volume = 1600 mm³) and right fornix (56,81,57; volume = 1400 mm³). Whole-brain seeding was used, with 1,000,000 seeds placed. The anisotropy and angular thresholds used for tractography were selected based on pilot testing and anatomical plausibility. We used a curvature threshold of 45° and an anisotropy threshold of 0.15, consistent with prior studies of fornix structure. Step size was randomly chosen between 0.5 and 1.5 voxels, and fiber tracking excluded tracts shorter than 20 mm or longer than 300 mm. ROI placement was based on standard MNI coordinates and manually verified using subject-specific anatomical landmarks.

### Data Analysis

All statistical analyses were conducted using RStudio. Relationships between fornix integrity and task performance were assessed by comparing percentage correct and reaction times from the working memory (WM) and recognition tasks with diffusion tensor imaging (DTI) metrics of the fornix.

Independent t-tests were performed to examine sex differences in each DTI metric and to compare measurements between the left and right fornix. Pearson’s correlations were calculated between each DTI metric and: (1) the change in reaction time between the first and third WM runs, (2) percentage correct in the recognition task, and (3) reaction times for old and new images in the recognition task.

To evaluate the interaction between fornix integrity and age, multivariate regression was used to assess the additional contribution of age to the model.

## Results

Initial analyses revealed no significant effects of sex on any behavioral or neuroimaging metrics, including reaction time, accuracy, or tractography measures. Similarly, working memory performance did not differ significantly across participants and was not associated with any structural indices examined in this study. Given the absence of meaningful variation in these domains, subsequent analyses focused on recognition memory performance, where robust structure-behavioral relationships emerged.

### Whole-Brain Fixel-Based Results

Fixel-based analysis revealed both dissociable and convergent patterns across FD, log(FC), and FDC in relation to aging and recognition memory speed. For FD, only the contrast testing the quadratic interaction of age and reaction time (Contrast 5) revealed significant effects. Significant clusters were localized to the left superior longitudinal fasciculus, a fronto-parietal association pathway situated lateral to the corona radiata and arching over the insula. Given its role in integrating attention, working memory, and cognitive control processes, structural variation in this tract may reflect individual differences in higher-order cognitive performance across development (Figure 1). This pattern suggests microstructural preservation or adaptive remodeling in attention-related pathways that support the capacity to sustain effort in memory retrieval tasks.

**Figure 1.**
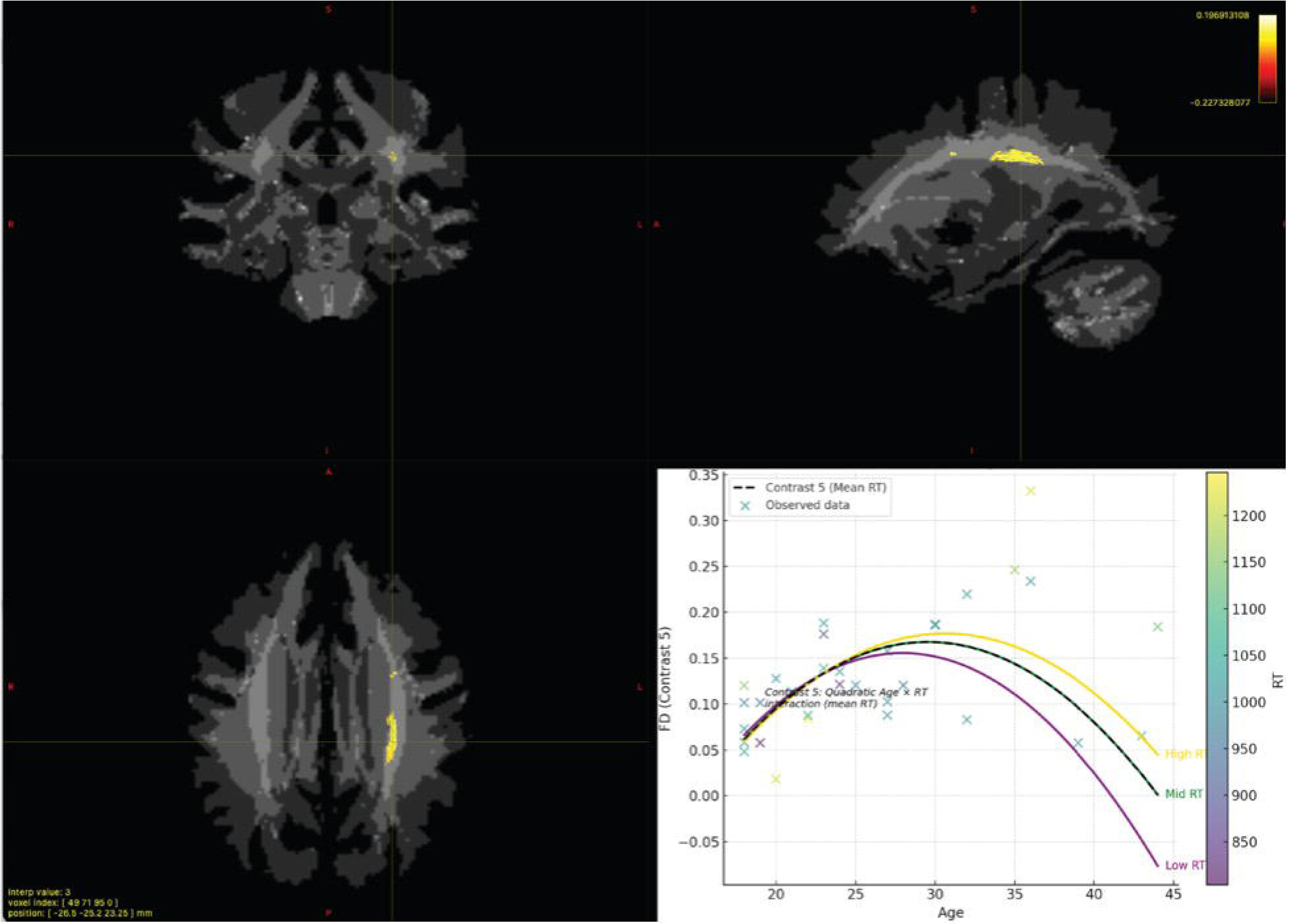
FD Contrast 5. Fiber density (FD) increased in older adults with slower reaction times (RT), with significant effects localized to the left superior longitudinal fasciculus—a frontoparietal association tract lateral to the corona radiata. This reflected a significant quadratic Age × RT interaction (b = 2.72×10⁻⁵), extracted from fixel-based analysis within clusters surviving FWE correction at p < .05 (R² = 0.276). These results suggest microstructural remodeling in aging as adaptive for cognitive control and behavioral inhibition.

In contrast, fiber cross-section (log(FC)) exhibited broader and more diverse effects. The most robust finding was the quadratic interaction of age and reaction time (Contrast 5), which revealed increased cross-sectional area in the anterior body and genu of the corpus callosum, a major interhemispheric white matter pathway connecting homologous regions of the prefrontal cortex (Figure 2). This tract supports cognitive control and executive functions by facilitating communication between the frontal lobes, and the observed pattern may reflect a greater capacity for cognitive control and behavioral inhibition. In this case, that meant not pushing one of the response buttons until was more certain about the correct answer. A significant linear interaction between age and reaction time (Contrast 1), although statistically weaker, revealed divergent slopes in anterior white matter, consistent with behaviorally modulated patterns of structural maturation and age-related decline (Supplementary Figure S1).

**Figure 2.**
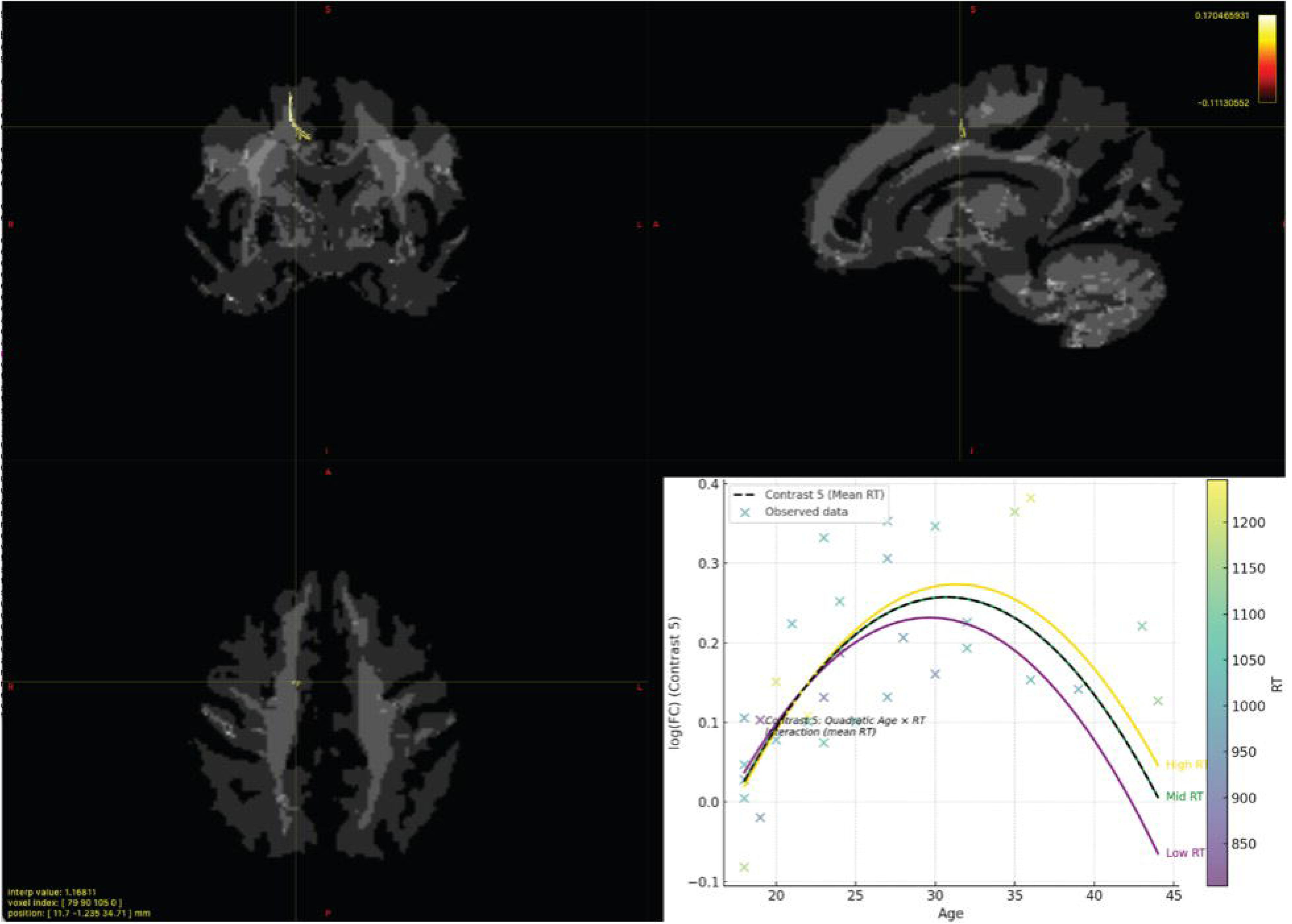
log(FC) Contrast 5. Nonlinear age-related increases in log-transformed fiber cross-section (log(FC)) were most pronounced among slower responders, with significant effects centered in the anterior body and genu of the corpus callosum, a key interhemispheric pathways linking bilateral prefrontal cortices. The model identified significant terms including Age² (b = 2.48×10⁻⁵), Age × RT (b = 2.15×10⁻⁵), and Age² × RT (b = –2.52×10⁻⁵) within FWE-corrected fixels (R² = 0.317). These findings support the idea of compensatory tract enlargement in response to age-related performance decline.

Additionally, two contrasts examining age-stratified RT effects identified modulated developmental trajectories: Contrast 9 highlighted enhanced maturation in log(FC) among slower- performing younger adults (Supplementary Figure S2), while Contrast 12 revealed a nonlinear (inverted-U) age effect, with peak cross-sectional area in midlife followed by a decline (Supplementary Figure S3).

FDC results closely paralleled those observed for log(FC), reinforcing the role of macrostructural change in these interactions. Contrast 5 revealed increased FDC among slower older adults in the splenium of the corpus callosum, with extension into the forceps major and adjacent posterior cingulum bundle (Figure 3). These posterior interhemispheric and limbic pathways are involved in visuospatial integration and memory processing, suggesting that greater cognitive control in these domains may reflect micro- and macrostructural maturation and preservation that supports improved accuracy. Similar patterns were observed in Contrast 9, where slower individuals exhibited preserved or enhanced FDC in left-lateralized regions (Supplementary Figure S4), and in Contrast 12, which revealed a nonlinear trajectory of FDC across the adult lifespan (Supplementary Figure S5).

**Figure 3.**
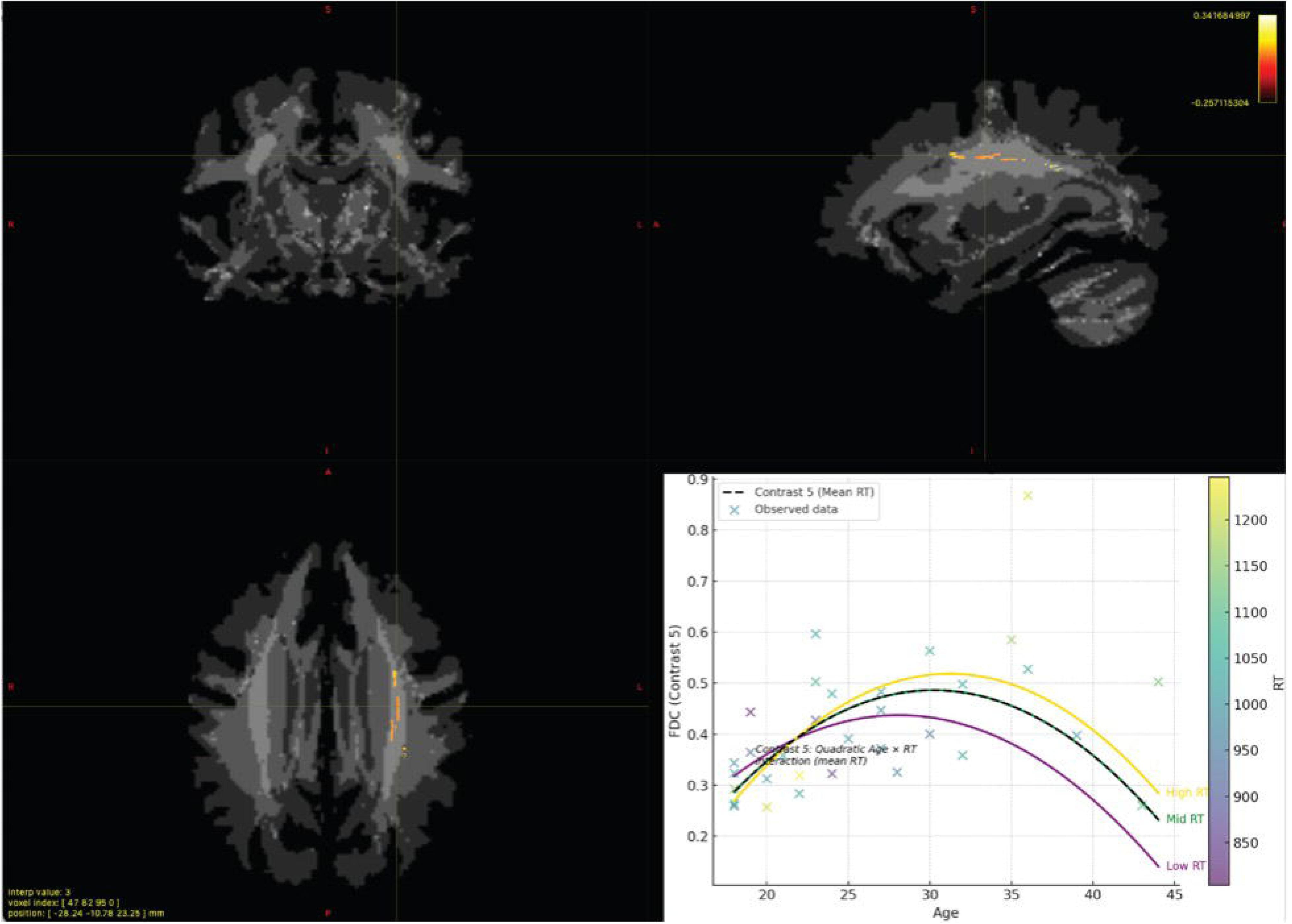
FDC Contrast 5. FDC demonstrated a significant quadratic Age × RT interaction similar to log(FC), with increased FDC among older adults with slower responses. Significant effects were localized to the splenium of the corpus callosum, with extensions into the forceps major and adjacent posterior cingulum bundle, a posterior white matter pathways implicated in visuospatial integration and memory. Significant terms included Age² (b = 2.45×10⁻⁵), Age × RT (b = 2.22×10⁻⁵), and Age² × RT (b = –2.46×10⁻⁵), all observed in p < .05 FWE-corrected clusters (R² = 0.345). This emphasizes the joint contribution of micro- and macrostructural adaptation to compensatory white matter organization in aging.

Together, these fixel-based findings highlight the left superior longitudinal fasciculus as a core region exhibiting sensitivity to age, performance, and their interaction. While fiber density changes were more focal, log(FC) and FDC captured broader, performance-dependent adaptations, supporting the notion that frontoparietal white matter structure reflects both a protracted period of maturation, peaking in midlife, and beginning to decline in older age.

### Fornix Tractography Results

There were no statistically significant sex differences in any fornix tractography metrics. Diffusion MRI tractography analyses were conducted separately for left and right fornix metrics, as well as for combined values, in relation to performance and reaction times. Comparisons between left and right fornix diffusion metrics revealed no significant differences, with the exception of tract count. A paired-samples *t*-test indicated that the number of streamlines was significantly greater in the left fornix compared to the right, *t*(31) = 2.36, *p* = .024. Six participants exhibited higher tract counts on the right, while the remainder showed a leftward asymmetry (see Table S1).

To increase statistical power, Groups A and B were combined for tractography-behavior correlations. In the combined group, fornix fractional anisotropy (FA) was significantly negatively correlated with RT for new items in the LTM task (*r* = −.41, *p* = .035; see Figure 4). In addition, RT was positively correlated with accuracy scores (*r* = .40, *p* = .040; see Figure 5), indicating that slower responses were associated with greater accuracy overall.

**Figure 4.**
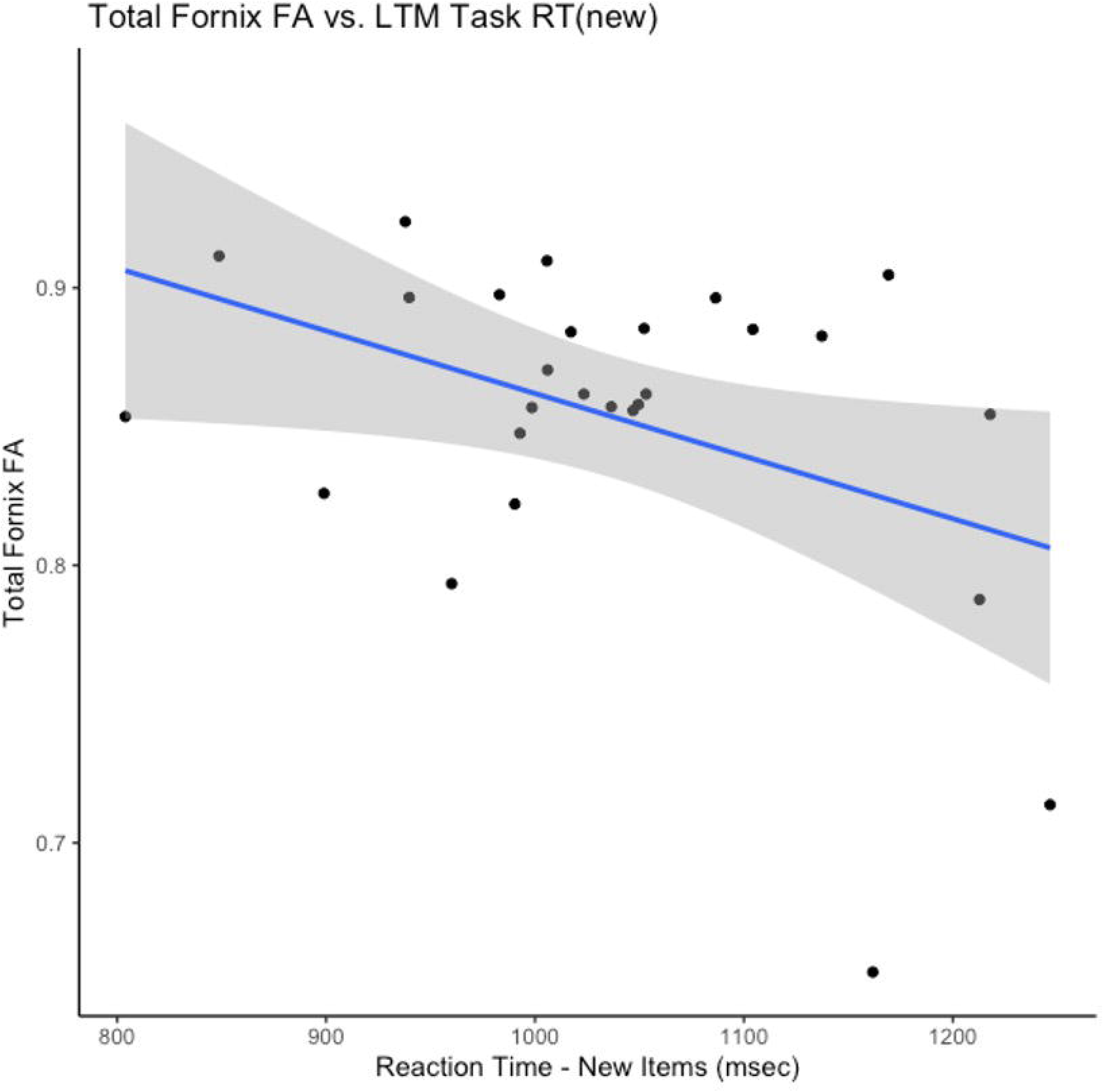
Fornix FA is negatively associated with reaction time to novel stimuli. Scatterplot showing the significant negative correlation between fractional anisotropy (FA) of the fornix and reaction time (RT) to new items in the long-term memory (LTM) task (*r* = −.41, *p* = .035). Greater fornix integrity was associated with faster recognition responses, suggesting that white matter microstructure contributes to increased speed of response.

**Figure 5.**
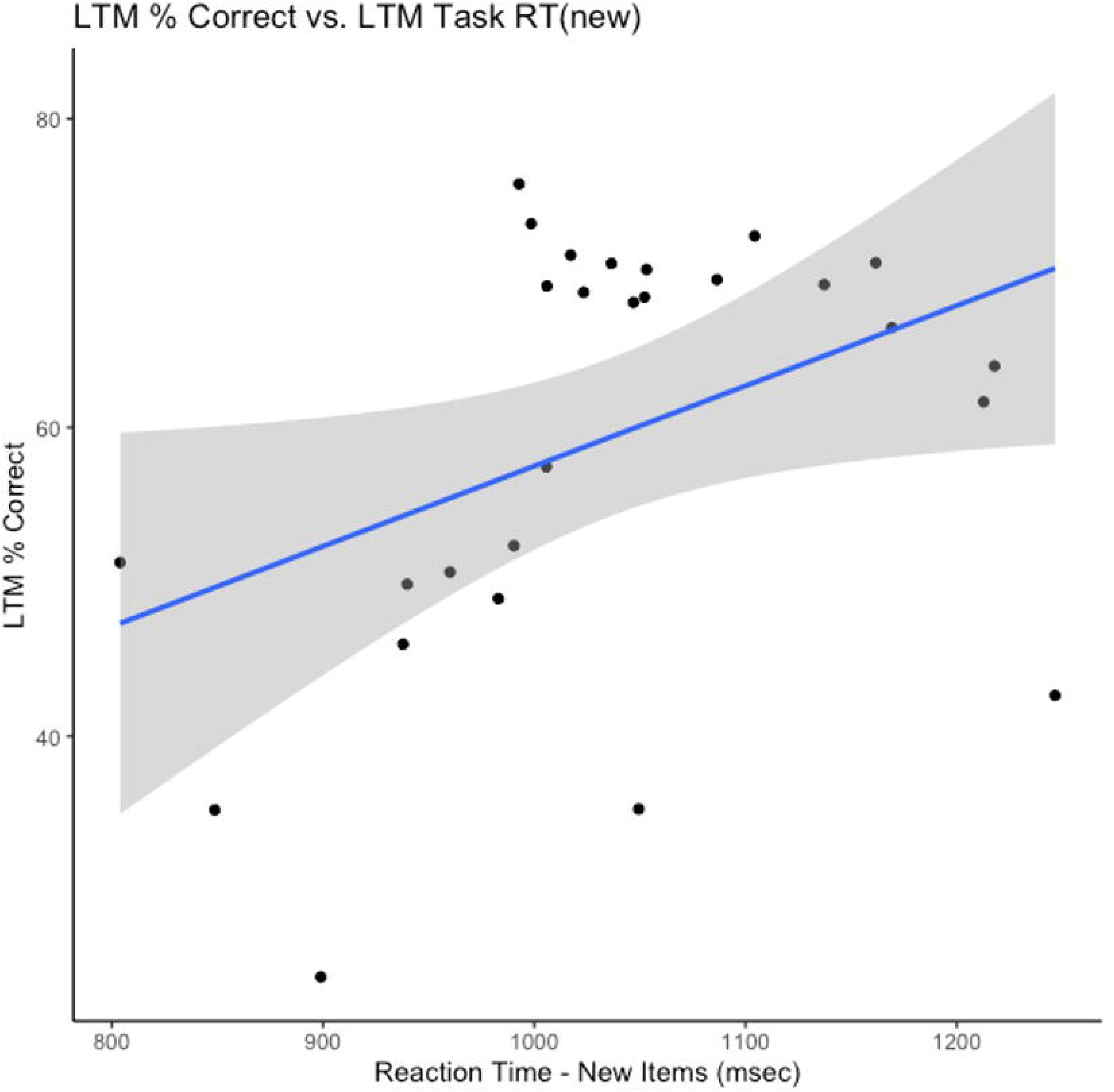
Slower reaction times are associated with increased accuracy in recognition memory. Scatterplot showing the significant positive correlation between RT to new items and LTM accuracy scores (*r* = .40, *p* = .040). Participants who responded more slowly tended to perform more accurately, demonstrating a trade-off between speed and accuracy (Reed, 1973).

A multiple linear regression model including both FA and age deviation as predictors of RT was statistically significant, *F*(2, 24) = 6.08, *p* = .007, explaining 34% of the variance (*R²* = .34, adjusted *R²* = .28). Both predictors contributed uniquely to the model: greater age deviation was associated with slower RTs (β = 6.13, *SE* = 2.47, *t* = 2.49, *p* = .020), while higher fornix FA predicted faster RTs (β = −973.63, *SE* = 315.04, *t* = −3.09, *p* = .005). The association illustrated in Figure 6 is both statistically and biologically meaningful. For example, a 0.05 increase in FA corresponds to an estimated 48.68 ms reduction in RT, a magnitude consistent with known white matter contributions to processing speed (Kerchner et al., 2012; Madden et al., 2004; Revie & Metzler-Baddeley, 2024).

**Figure 6.**
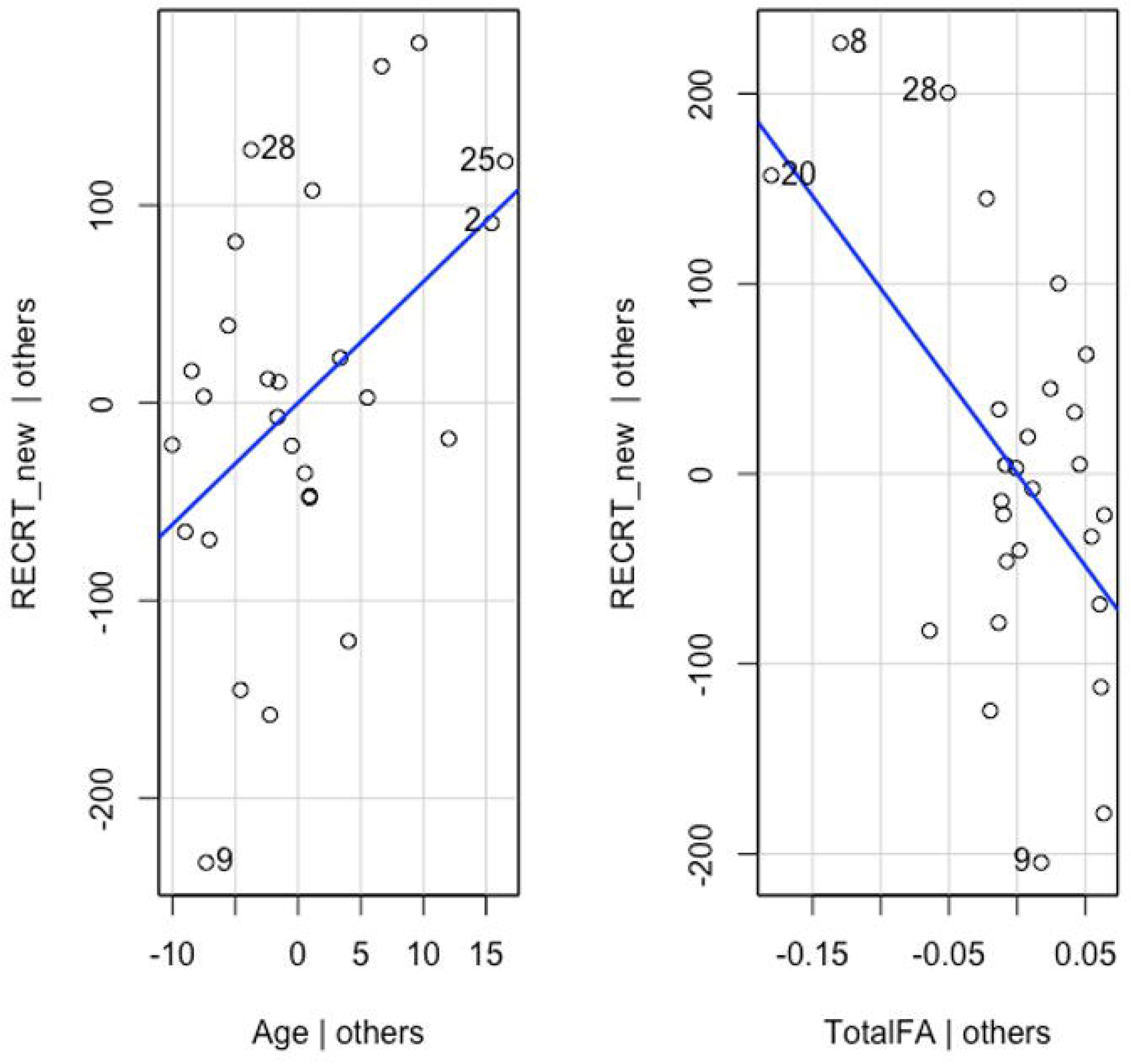
Fornix FA and age deviation jointly predict reaction time in memory retrieval. Partial regression plots show unique contributions of age deviation (left panel) and fornix FA (right panel) to RT in a multiple linear regression model (*F*(2, 24) = 6.08, *p* = .007, *R²* = .34), with age positively predicting RT (β = 6.13, *p* = .020) and FA negatively predicting RT (β = −973.63, *p* = .005). This model demonstrates that both age and fornix microstructure independently contribute to variability in recognition speed.

## Discussion

The main findings from the whole-brain fixel-based analyses provide converging evidence that age-related differences in white matter structure are both behaviorally modulated and regionally specific. Across primary contrasts, individuals with slower recognition memory responses exhibited greater fiber density (FD), fiber cross-section (log(FC)), and the combined metric of fiber density and cross-section (FDC) in frontoparietal and callosal tracts, particularly among older adults. FD effects were especially prominent in the left superior longitudinal fasciculus, a fronto- parietal association pathway linked to attention and cognitive control. These effects were strongest in regions supporting attentional and integrative processing, with log(FC) and FDC showing heightened sensitivity to quadratic interactions between age and performance. For log(FC), significant effects were centered in the anterior body and genu of the corpus callosum, which connect homologous prefrontal regions and may support enhanced interhemispheric communication and behavioral inhibition. Memory processing for the complex visual stimuli used in this study has previously been documented to elicit functional interhemispheric communication as measured using EEG (Plaska et al., 2022). This pattern suggests ongoing structural development during early and middle adulthood that supports greater retrieval accuracy and behavioral regulation. These patterns align with models of adaptive neuroplasticity (Bennett & Madden, 2014; Fjell & Walhovd, 2010) and are consistent with evidence that aerobic fitness and cognitive engagement can bolster white matter integrity in aging (Hayes et al., 2015; Lee et al., 2024). Notably, Bendlin et al. (Bendlin et al., 2010) also observed that white matter microstructure (e.g., FA and MD), rather than volume alone, shows nonlinear age-related changes and is tightly linked to cognitive performance, supporting our interpretation that micro- and macrostructural remodeling are functionally meaningful.

Supplementary subgroup contrasts revealed that performance-related white matter differences emerge during early adulthood, while older adults exhibited inverted-U trajectories in log(FC) and FDC. These patterns mirror findings from lifespan imaging studies showing age-specific structural adaptations in cortical and white matter networks and reinforce the view that white matter aging is dynamic and nonlinear (Conte et al., 2024). Bendlin et al. (Bendlin et al., 2010) similarly found that microstructural changes begin earlier in life and accelerate with age, even in neurologically healthy adults, providing critical support for our lifespan-informed subgroup approach. Variation in these trajectories may reflect cognitive reserve, baseline structural health, or cognitive lifestyle, all factors shown to shape neural plasticity and transfer in training contexts (Basak et al., 2020; Lee et al., 2024).

Taken together, the fixel-based results support a model of regionally specific, behaviorally responsive plasticity in aging. Increased fiber cross-section or density in some tracts may reflect adaptive remodeling, especially in higher-order association pathways. FDC effects extended into the splenium of the corpus callosum, forceps major, and adjacent posterior cingulum bundle - tracts that support visuospatial integration and memory processing - suggesting that older adults with slower RTs may recruit both anterior and posterior networks in response to increased cognitive demands. Parallel effects across log(FC) and FDC underscore the relevance of macrostructural remodeling - not just microstructural loss - as a key feature of aging. The use of whole-brain FBA with rigorous family-wise error correction strengthens confidence in these effects By linking individual performance differences to directionally meaningful, tract-specific structural changes, our findings advance models of cognitive aging and underscore the value of lifespan- informed, multimodal approaches (Charlton et al., 2010; Penke et al., 2010).

Complementing these whole-brain findings, hypothesis-driven analysis of the fornix - a key hippocampal efferent - demonstrated that greater fornix FA was associated with faster reaction times, independent of age. Both age and fornix FA were significant predictors in a regression model accounting for 34% of the variance in retrieval speed. This suggests that faster responses in memory circuits may rely on preserved microstructure rather than compensatory changes. Prior studies highlight the fornix’s role in hippocampal–diencephalic connectivity and memory consolidation (Aggleton et al., 2000; Rudebeck et al., 2009), and show that fornix FA predicts episodic memory performance and early cognitive decline (Fletcher et al., 2013; Metzler-Baddeley et al., 2011; Metzler-Baddeley et al., 2012; Mielke et al., 2012). Our findings suggest that fornix integrity contributes to retrieval speed in visual memory, whereas accuracy may depend on broader white matter architecture. Because the task required prompt responses to register, participants who were both fast enough to be counted and slow enough to be accurate may reflect peak white matter maturation. This implies an optimal window of structural development prior to age-related decline that enables a balance between speed and accuracy in memory tasks. This echoes Bendlin’s conclusion that microstructural integrity better explains cognitive function in aging than gross volumetric measures, highlighting the value of high-resolution diffusion metrics.

Importantly, the fornix findings diverge from the macrostructural expansion seen in frontoparietal association tracts. Here, higher FA was linked to faster responses without concurrent morphological enlargement, suggesting that microstructural preservation, not remodeling, supports speed in this circuit. Variability in fornix integrity may also influence responsiveness to cognitive interventions, as prior work shows that structural health moderates memory training outcomes (Basak et al., 2020).

While fornix FA and age together explained a substantial portion of the variance in RT, considerable variability remained. This implies other factors, such as attentional control, prefrontal function, or large-scale connectivity, likely contribute to performance differences (Braver & West, 2011; Nyberg et al., 2012). Functional imaging studies suggest that training-related changes in network efficiency and inter-regional communication may further influence retrieval speed (Andrushko et al., 2023; Dresler et al., 2017).

Several limitations warrant consideration. First, the cross-sectional design limits causal inference regarding age-related change or its effect on memory. The age range is limited to early and middle adulthood using a small, limited sample. Longitudinal studies and larger samples are needed to distinguish true developmental trajectories from cohort effects. Second, although FBA improves specificity over tensor-based methods, it may lack sensitivity in detecting effects within small or complex tracts like the fornix. Its narrow structure, curvature, and proximity to CSF increase vulnerability to partial volume effects and registration errors, which may explain its absence in whole-brain results despite robust ROI findings. Third, although fornix tractography was hypothesis-driven and anatomically grounded, the lack of multiple comparison correction raises concern for inflated Type I error. However, consistency with prior studies helps mitigate this risk. Finally, the recognition task emphasized speed over accuracy and did not differentiate recollection from familiarity, limiting our ability to map structural features onto specific mnemonic processes. Future work should employ tasks that dissociate retrieval subprocesses and include complementary accuracy metrics.

Future research should incorporate longitudinal, multimodal imaging to trace structural adaptation over time and identify interventions that preserve or enhance white matter integrity in aging. Targeted studies of hippocampal subfields, network connectivity, and individual responsiveness to cognitive training will be critical to refining models of resilience and personalizing strategies to support healthy cognitive aging.

