## Supplementary Information for "Linking White Matter Integrity to Recognition Memory Speed: Fixel-Based and Fornix Analyses in Young to Middle Adulthood"

**Supplemental Information**

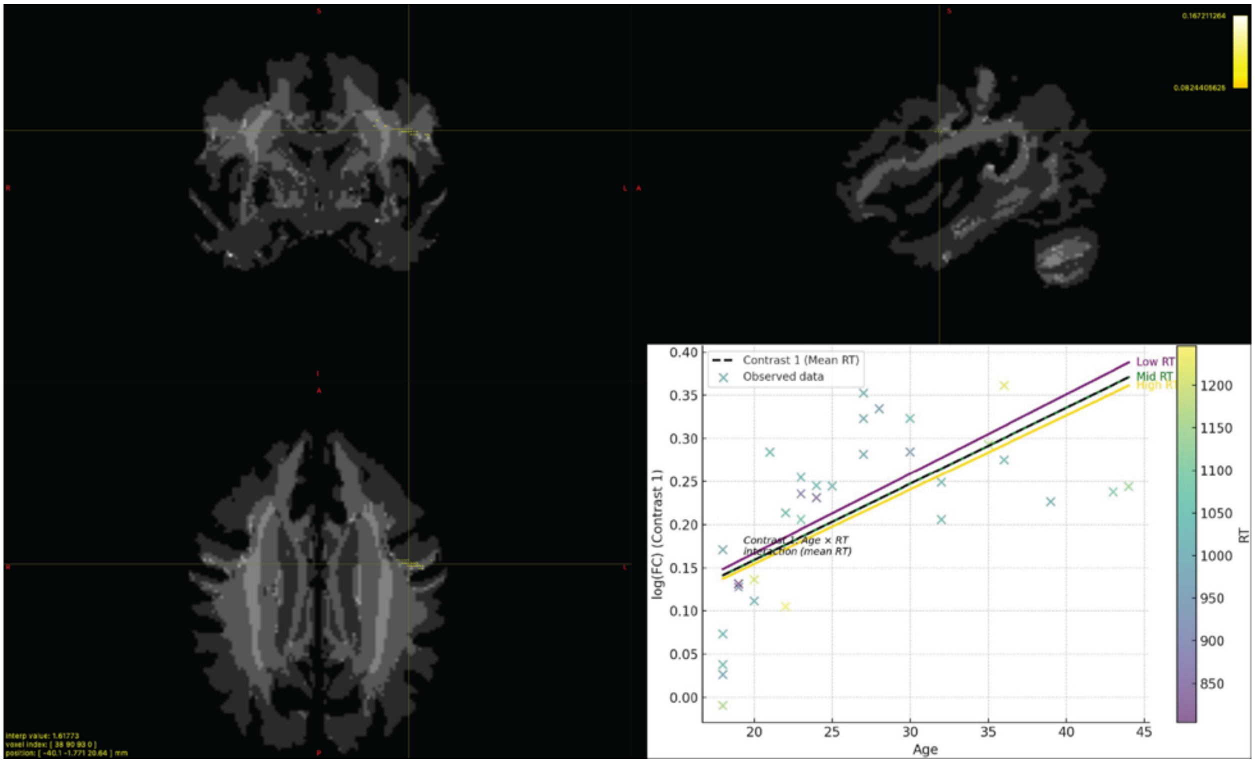


**Supplemental Figure S1 log(FC) Contrast 1.** In fixels surviving p < 0.05 FWE correction, age-related increases in log-transformed fiber cross-section (log(FC)) were evident across all response time (RT) groups. However, individuals with faster responses (low RT) exhibited a steeper age-related increase in log(FC) compared to slower responders (high RT), whose trajectories were more attenuated. This pattern suggests that cognitive performance may modulate the extent of age-related macrostructural plasticity in anterior white matter.

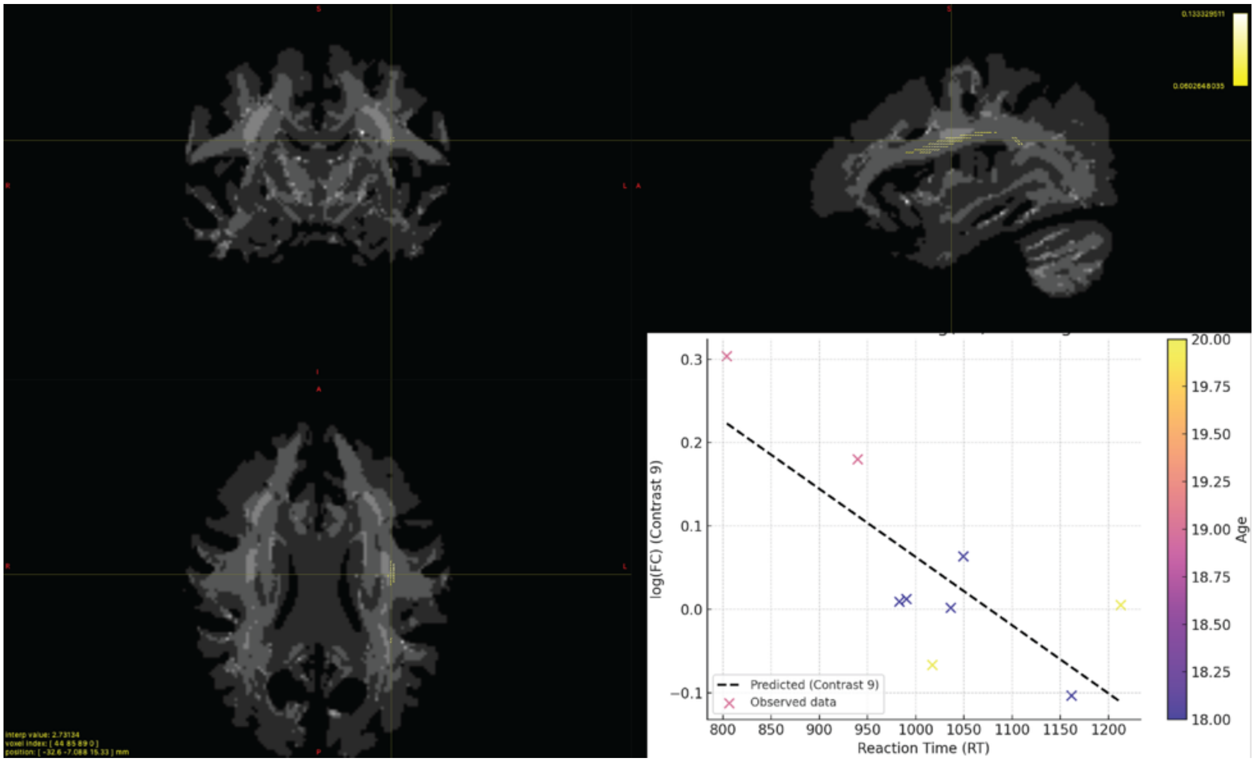


**Supplemental Figure S2**. **log(FC) Contrast 9.** Among participants in the youngest age quartile, faster RT was associated with larger log-transformed fiber cross-section (log(FC)) in the left superior longitudinal fasciculus (SLF). These effects, identified in fixels surviving p < 0.05 FWE correction, suggest that performance-related variation in white matter morphology may be detectable even in early adulthood.

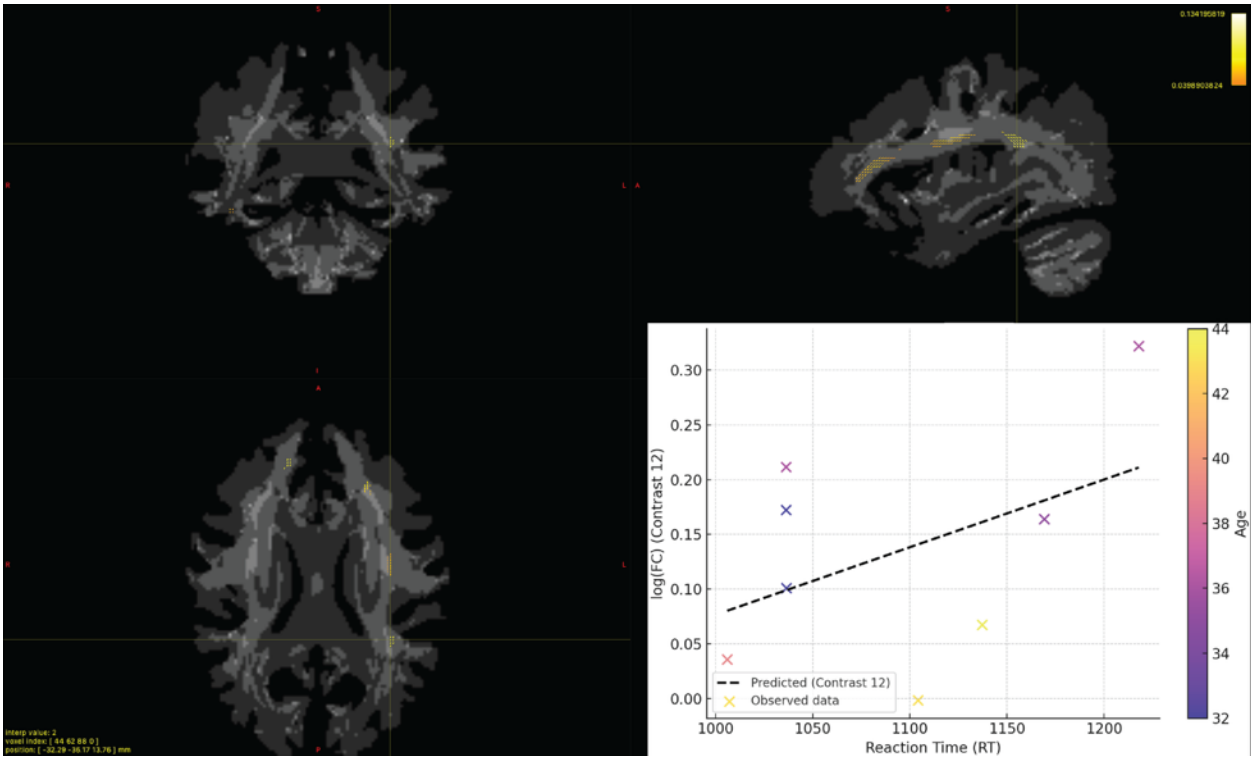


**Supplemental Figure S3. log(FC) Contrast 12.** Among participants in the highest age quartile, fixels passing the $p < 0.05$ FWE threshold showed a general pattern of increased log(FC) among slower individuals, pointing to possible structural compensation or maintenance of fiber bundle size in later life.

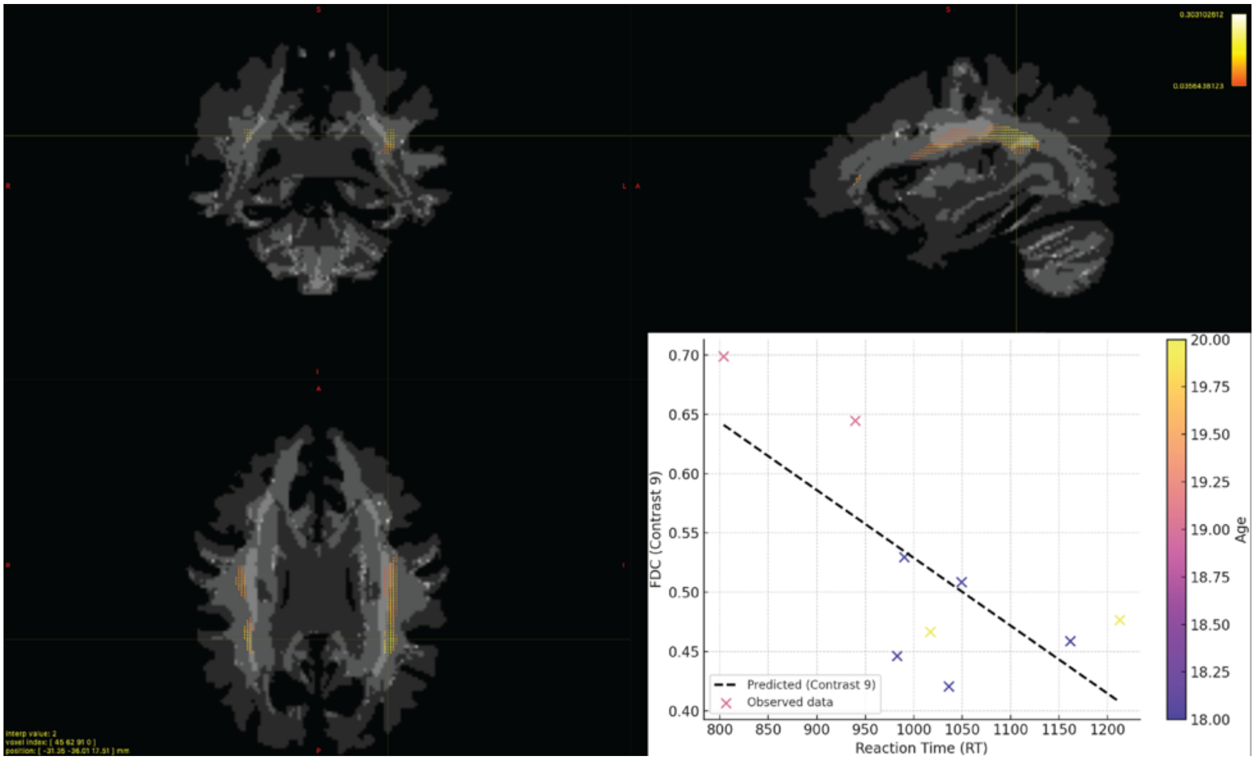


**Supplemental Figure S4. FDC Contrast 9.** In fixels within the left and to a lesser extent right superior longitudinal fasciculus (SLF) that survived $p < 0.05$ FWE correction, slower responders in the youngest age quartile exhibited modest decreases in fiber density and cross-section (FDC). This trend suggests that individual differences in retrieval speed may be associated with subtle reductions in white matter structure even in early adulthood.

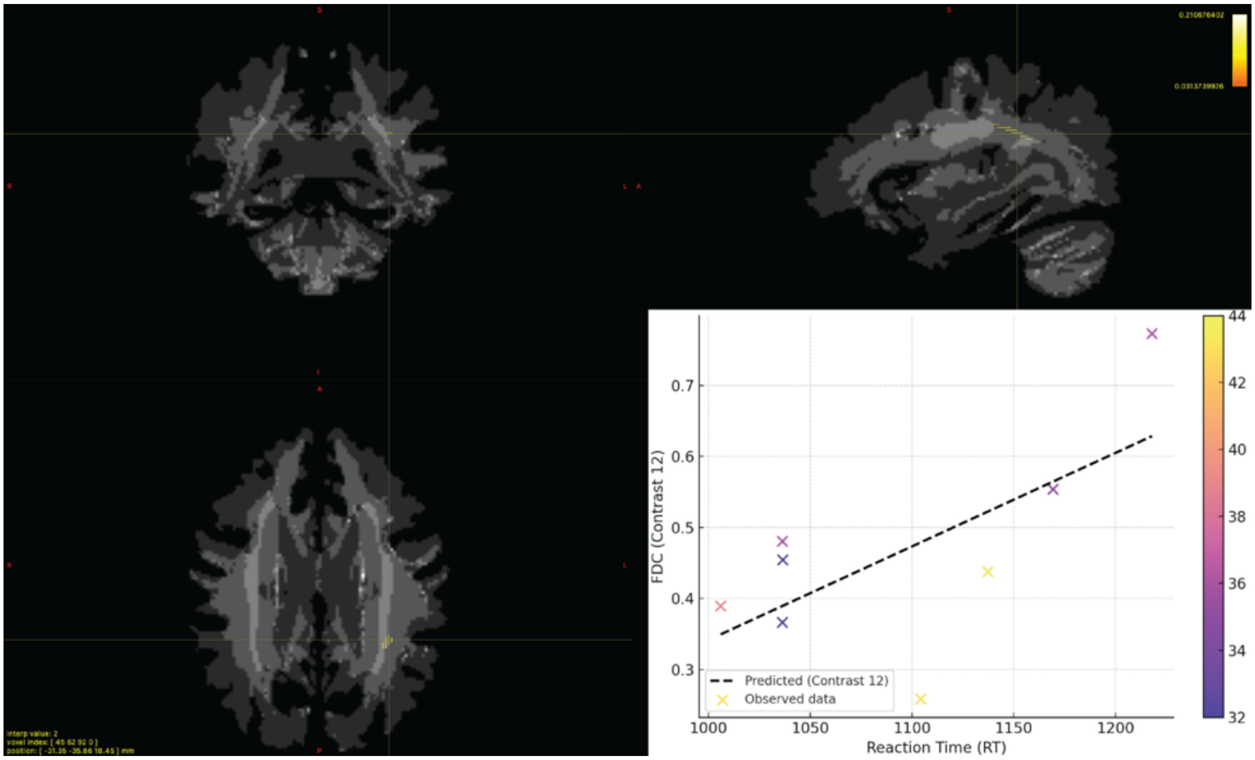


**Supplemental Figure S5. FDC Contrast 12.** Among participants in the highest age quartile, slower responders exhibited greater FDC in left posterior fixels meeting the $p < 0.05$ FWE threshold. This pattern suggests that combined micro- and macrostructural integrity may be preserved or enhanced in response to reduced processing speed in aging.

| Group | Subject ID | # of Tracts (Left) | # of Tracts (Right) | Difference Score | Total FA |
|---|---|---|---|---|---|
| A | SUJ_36 | 11516 | 10569 | 947 | 0.86 |
| A | SUJ_40 | 12991 | 14461 | -1470 | 0.89 |
| A | SUJ_42 | 10535 | 11697 | -1162 | 0.79 |
| A | SUJ_46 | 16825 | 12665 | 4160 | 0.9 |
| A | SUJ_47 | 7556 | 7648 | -92 | 0.83 |
| A | SUJ_48 | 12667 | 11417 | 1250 | 0.86 |
| A | SUJ_49 | 14784 | 12908 | 1876 | 0.92 |
| A | SUJ_50 | 9193 | 8807 | 386 | 0.71 |
| A | SUJ_51 | 15929 | 11073 | 4856 | 0.85 |
| A | SUJ_57 | 18055 | 15533 | 2522 | 0.9 |
| A | SUJ_59 | 11604 | 12644 | -1040 | 0.91 |
| A | SUJ_60 | 15388 | 14946 | 442 | 0.91 |
| A | SUJ_64 | 14965 | 12375 | 2590 | 0.9 |
| A | SUJ_65 | 7533 | 7749 | -216 | 0.82 |
| A | SUJ_66 | 8922 | 7498 | 1424 | 0.85 |
| A | SUJ_67 | 10454 | 10130 | 324 | 0.89 |
| B | SUJ_68 | 10757 | 8232 | 2525 | 0.86 |
| B | SUJ_69 | 12214 | 9431 | 2783 | 0.85 |
| B | SUJ_70 | 22520 | 14019 | 8501 | 0.89 |
| B | SUJ_71 | 10629 | 9021 | 1608 | 0.65 |
| B | SUJ_72 | 10479 | 9677 | 802 | 0.90 |
| B | SUJ_73 | 15347 | 10623 | 4724 | 0.87 |
| B | SUJ_74 | 13056 | 11086 | 1970 | 0.86 |
| B | SUJ_75 | 12940 | 13333 | -393 | 0.90 |
| B | SUJ_76 | 10341 | 8462 | 1879 | 0.88 |
| B | SUJ_77 | 17554 | 15237 | 2317 | 0.86 |
| B | SUJ_78 | 12117 | 8073 | 4044 | 0.96 |
| B | SUJ_79 | 12989 | 10212 | 2777 | 0.79 |
| B | SUJ_80 | 13636 | 12658 | 978 | 0.86 |
| B | SUJ_81 | 11519 | 9569 | 1950 | 0.97 |
| B | SUJ_82 | 14857 | 11470 | 3387 | 0.88 |
| B | SUJ_83 | 11805 | 9373 | 2432 | 0.86 |

**Supplemental Table S1.** Individual subject data for fornix tractography. The number of tracts associated with the left and right fornix and the difference score (DF) between them (left- right= DF), followed by the total FA of left and right fornixes combined.
